# Artemisinin acts by inhibiting *Plasmodium falciparum* Ddi1, a retropepsin, resulting into the accumulation of ubiquitinated proteins

**DOI:** 10.1101/2021.07.12.452004

**Authors:** Noah Machuki Onchieku, Sonam Kumari, Rajan Pandey, Vaibhav Sharma, Mohit Kumar, Arunaditya Deshmukh, Inderjeet Kaur, Asif Mohmmed, Dinesh Gupta, Daniel Kiboi, Naseem Gaur, Pawan Malhotra

**Affiliations:** Malaria Biology Group, International Centre for Genetic Engineering, and Biotechnology, Aruna Asaf Ali Marg, New Delhi, India; Department of Biochemistry, Jomo Kenyatta University of Agriculture and Technology, Nairobi, Kenya; Yeast Biofuel Group, International Centre for Genetic Engineering and Biotechnology, Aruna Asaf Ali Marg, New Delhi, India; Translational Bioinformatics Group, International Centre for Genetic Engineering and Biotechnology, Aruna Asaf Ali Marg, New Delhi, India; Parasite Cell Biology Group, International Centre for Genetic Engineering, and Biotechnology, Aruna Asaf Ali Marg, New Delhi, India

**Author notes:** Corresponding author: Pawan Malhotra.

**Keywords:** Artemisinin, *Plasmodium falciparum*, DNA Damage, Ddi1, Ubiquitin-Proteasome Pathway

## Abstract

Reduced sensitivity of the human malaria parasite, *Plasmodium falciparum,* to Artemisinin and its derivatives (ARTs) threatens the global efforts towards eliminating malaria. ARTs have been shown to cause ubiquitous cellular and genetic insults, which results in the activation of the unfolded protein response (UPR) pathways. The UPR restores protein homeostasis, which otherwise would be toxic to cellular survival. Here, we interrogated the role of DNA-damage inducible protein 1 (*Pf*Ddi1), a unique proteasome-interacting retropepsin in mediating the actions of the ARTs. We demonstrate that *Pf*Ddi1 is an active A_2_ family protease that hydrolyzes ubiquitinated substrates. We further show that treatment with ARTs lead to the accumulation of ubiquitinated proteins in the parasites and blocks the destruction of the ubiquitinated substrates by *Pf*Ddi1. Besides, whereas the *Pf*Ddi1 is predominantly localised in the cytoplasm, exposure of the parasites to ARTs leads to DNA fragmentation and increased recruitment of the *Pf*Ddi1 into the nucleus. Furthermore, Ddi1 knock-out *Saccharomyces cerevisiae* cells are more suceptible to ARTs and the *Pf*DdI1 protein robustly restores the corresponding functions in the knock-out cells. Together, these results show that ARTs act by inducing DNA and protein damage, and impairing the damage recovery by inhibiting the activity of *Pf*Ddi1, an essential ubiquitin-proteasome retropepsin.

## Introduction

Artemisinin and its derivatives (ARTs) are components of mainstay drugs for the treatment of malaria caused by the *Plasmodium falciparum* parasite^1^. However, the emergence and spread of resistance towards the artemisinins poses an imminent danger towards the global efforts to eliminate malaria^2^. Historically, the spread of malaria drug resistance mechanisms from South East (SE) Asia to India is a crucial “stepping stone” to the eventual introduction in Africa^3,4^. Regrettably, recent evidence has shown the presence of Artemisinin resistant *P. falciparum* in India^5^. This situation does not only pose a grave danger to public health in these countries but also in sub-Saharan Africa, a continent most affected by malaria^1^. Whereas there is the most reliable evidence linking mutations in the Kelch domain protein (K13-propeller; PF3D7_1343700) with parasite tolerance to artemisinin^6^, insufficient knowledge on the molecular mechanisms of artemisinin action hampers definitive conclusion. Understanding the mechanisms of action and resistance of artemisinin, therefore, would not only provide a basis for identifying new targets but also be useful to the development of new alternative compounds that thwart and antagonize the emergence of resistance.

Recent reports have demonstrated the promiscuous nature of artemisinin-mediated cellular damages^**7–11**^. For instance, besides the ubiquitous protein insults^12^, Artemisinin has been attributed to DNA damage mediated by reactive oxygen species (ROS)^9^. Consequently, the damage would be expected to trigger stress response or unfolded protein response (UPR) pathways^13,14^, such as the ubiquitin-proteasome system (UPS)^12^. The UPS degrades unfolded/damaged proteins that would otherwise be toxic to cells. Interestingly, evidence has associated the K13-propeller protein with ubiquitination^8,15^, and inhibitors of the UPS have been shown to enhance the action of artemisinin against *P. falciparum* parasites^16–18^. Artemisinin inhibits the UPS and the changes to this system mediate parasite tolerance to artemisinin pressure^12,16,19^. However, molecular data on the role of the UPS in mediating the action/resistance of artemisin in *P. falciparum* parasites remains scanty.

*Pf*Ddi1, an essential retropepsin in the UPS^20,21^, has been shown to compensate for proteasome dysfunction and its knock out leads to polyubiquitination of proteins in both yeast and *Toxoplasma gondii* cells^22–24^. It is feasible, therefore, to speculate that artemisinin might be compromising the activity of *Pf*Ddi1 in restoring protein homeostasis following the damage. Here, we identify the *Pf*Ddi1, commonly referred to as the proteasome shuttle protein, and investigate its role in mediating the actions of artemisinin. Binding and enzymatic assays demonstrate that *Pf*Ddi1 is an active proteasome reptropepsin that cleaves ubiquitinated substrates. We show that artemisinin enhances polyubiquitination of parasite proteins and inhibits the activity of *Pf*Ddi1 in digesting the ubiquitinated proteins. In addition, the parasites’ exposure to artemisinin induces DNA fragmentation and increases recruitment of the *P*fDdi1 protein into the nucleus. Besides, using yeast complementation studies, we show that whereas *Pf*Ddi1 is dispensable in yeast, *Pf*Ddi1 defficient *S. cerevisiae* cells display more susceptibility to artemisisnin pressure. The expression of *Pf*Ddi1 restores the functions in the corresponding Ddi1-knock out yeast cells. Our work thus gives insights into the role of the *Pf*Ddi1 and validates it as a vulnarable protein that could be the basis for the development of new chemotherapies against the *P. falciparum* malaria.

## Results

### *Pf*DdI1 is an active A_2_ family protease that hydrolyzes polyubiquitin substrates

Whereas *P. falciparum* parasites express three proteasome interacting proteins (PIPs); *Pf*Ddi1, Rad23 and Dsk2, deletion of only *Pf*Ddi1 has been proven to be toxic to the cells, thus indespensable^20,21,25^. Compared to the other *Pf*PIPs, *Pf*Ddi1 harbors a unique retroviral-protease like (RVP) domain besides the conventional ubiquitin-like (UBL) domain. Despite being characterized in other organisms, Ddi1 remains poorly undestood in *Plasmodium* spp. To functionally characterize the role of the *Pf*Ddi1 if any, we cloned, and expressed a histidine-tagged full length *PfDdi1*gene (PF3D7_1409300) in Rosetta (DE3) cells. The expressed recombinant *Pf*Ddi1 protein was analyzed by both Coomassie staining and Western blotting with α-His antibodies. The *Pf*Ddi1 protein was then purified under non-denaturing conditions, and it showed two discrete bands of ~44kDa and ~34 kDa sizes on SDS PAGE, suggesting that the ~34 kDa band is probably a processed fragment of the *Pf*Ddi1 protein (Fig. 1a and Supplementary Fig. 1). To confirm whether the ~34kDa band is indeed a processed product of the intact *Pf*Ddi1 protein, we analysed both bands by LC-MS/MS. The proteome analysis showed that the peptides identified in the LC-MS/MS analysis for each of the fragments corresponded to the *Pf*Ddi1 protein and interestingly, they both had the aspartic catalytic signature motif (DSG) (Supplementary Fig. 2). The purified recombinant *Pf*Ddi1 protein was then used to raise antibodies in mice and rabbits. The specificity of the antibodies to detect native *Pf*Ddi1was assessed by Western blot using trophozoite-rich *P. falciparum* blood stage parasite lysate. The mice or rabbit anti-*Pf*Ddi1 antibodies stained a band of the size expected for *Pf*Ddi1 in *P.falciparum (*Fig. 1b). Since *Pf*Ddi1 possesses a retroviral-like protease (RVP) domain, we next assessed the pepsin/cathepsin D, retropepsin or proteasome activity of the purified recombinant *Pf*DdI1 protein using the Bz-RGFFP-MNA, DABCYL-Gaba-SQNYPIVQ-EDANS or Suc-LLVY-AMC substrates, respectively. Unlike the cathepsin D substrate, 2.0 μM of the enzyme hydrolyzed DABCYL-Gaba-SQNYPIVQ-EDANS or Suc-LLVY-AMC at pH 5.0. The enzyme was more active on the retropepsin substrate, with a catalytic efficiency of ~3.8 × 10^5^ M^−1^ s^−1^ (*K_m_* = 4.135 ± 0.280 μM), compared to the proteasome specific substrate, with an efficiency of ~8.0 × 10^4^ M^−1^ s^−1^ (*K_m_* = 21.85 ± 4.135 μM) (Fig. 1c). Due to its ability to hydrolyze proteasome substrates, coupled with previous evidence that Ddi1 compensates for proteasome dysfunction^22^, we hypothesized that the *Pf*Ddi1 might harbor the ability to degrade polyubiquitinated proteins/substrates. Polyubiquitination serves as a recognition signal for the proteasome. Our data showed that, incubation of K^48^-linked polyubiquitin substrate with 2.0 μM *Pf*Ddi1enzyme led to significant cleavage of the substrate (Fig. 1d). Together, these findings demonstrate that *Pf*Ddi1 is an active retroviral protease that hydrolyzes polyubiquitin/proteasome substrates.

**Fig. 1:**
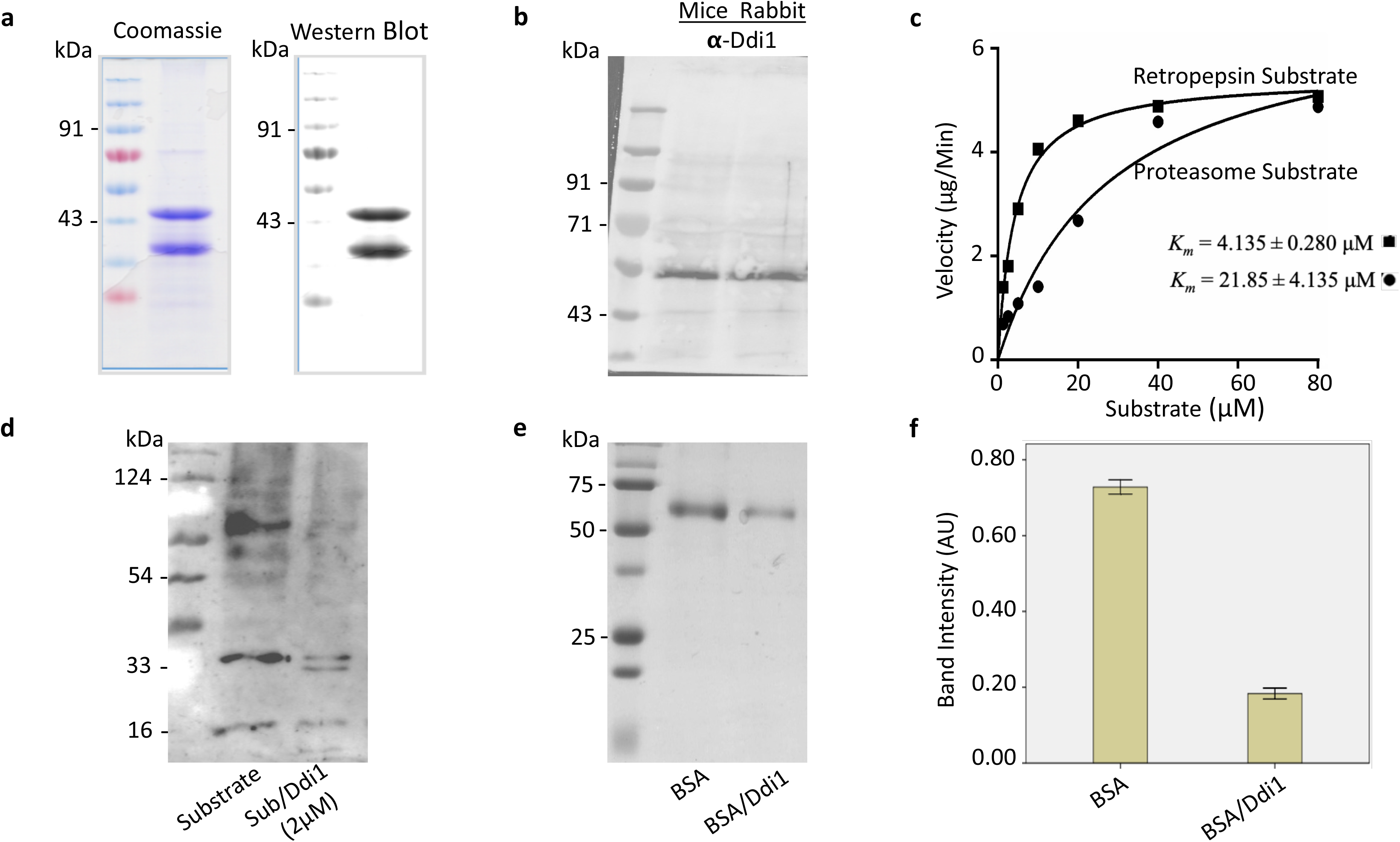
*Pf*Ddi1 hydrolyzes both peptide substrates and proteins. **(a)** Coomassie stained SDS-PAGE and Western blot analysis using α-His antibodies to detect the purified recombinant *Pf*Ddi1 (~44 kDa) and its processed fragment (~34 kDa). The soluble, recombinant protein was purified using the Ni-NTA resin. (b) Mice or rabbit ant-Ddi1 antibodies detected a band of ~49 kDa from a parasite lysate. (c) Kinetics of substrate hydrolysis. The enzyme was more active on the retropepsin (DABCYL-Gaba-SQNYPIVQ-EDANS) substrate compared to the proteasome (Suc-LLVY-AMC) substrate. (d) Western blot analysis showing the cleavage of polyubiquitin substrate by the *Pf*Ddi1 enzyme. The reaction was incubated for 1 hr at 37°C (e) Coomassie stained SDS-PAGE showing degradation of BSA by the recombinant *Pf*Ddi1. The samples were resolved in a 12% SDS-PAGE. (**f**) Quantification of the control and the degraded BSA band intensities (~66 kDa). The units are arbitrary (AU) and the bars show the mean **±** standard error for three independent reactions.

### *Pf*Ddi1 enzyme degrades Bovine serum albumin, BSA

Having demonstrated the ability of the recombinant *Pf*Ddi1 protein to hydrolyze peptide substrates, we assessed the capacity of the enzyme to degrade macromolecules. Compared with the control (BSA alone), incubation of BSA with the *Pf*Ddi1 protein resulted in the degradation of BSA, at pH 5.0. SDS-PAGE analysis of the test assay showed a significantly reduced BSA band intensity (~66kDa) (Fig. 1e and f). On the other hand, the *Pf*Ddi1 could not hydrolyze BSA at pH 7.0 (Supplementary Fig. 3). In *Leishmania major,* an acidic pH has been shown to be more favourable to the Ddi1 activity.^26^

### Artemisinin increases polyubiquitination in *P. falciparum* and blocks the activity of *Pf*DdI1 in degrading the polyubiquitinated substrates

Artemisinin has been shown to cause widespread damages to parasite proteins^7,8,11^. The damage invokes the unfolded protein response pathways as a means of tidying up. Here, we assessed the impact of artemisinin on global protein ubiquitination as well as on the activities of the *Pf*Ddi1 enzyme. Exposure of trophozoite-rich 3D7 *P. falciparum* parasites to 1.0 μM of artemisinin (a physiologically relevant dose^27^) for 2 hours resulted into accumulation of polyubiquitinated proteins. Similarly, Dihydroartemisinin (DHA; 1.0 μM, a biologically active ART metabolite) and MMS (0.05%) led to enhanced polyubiquitination, but not Lopinavir (50 μM) (Fig. 2a). The rapid protein polyubiquitination under artemisinin pressure invoked thoughts about its potential inhibition ability against the *Pf*Ddi1 enzyme activities. Indeed, artemisinin and its derivative, DHA, significantly inhibited the ability of *Pf*Ddi1 to degrade the polyubiquitinated substrate. Besides, artemisinin significantly inhibited the activity of *Pf*Ddi1 with both the retropepsin (71.4%) and proteasome (65.9%) specific substrates, as well as with BSA (Fig. 2b-f). Lopinavir (50 μM), a known HIV protease inhibitor, produced about 23% inhibition. Suprisingly, whereas MMS (0.05%) led to increased polyubiquitination, it enhanced the activities of *Pf*Ddi1 proteins against all the substrates (Fig. 2b-f). These data demonsrates the dual mechanisms of action of the artemisinins; by causing protein damage in the parasite and blocking tidying up by inhibiting the activities of *Pf*Ddi1 enzyme.

**Fig. 2:**
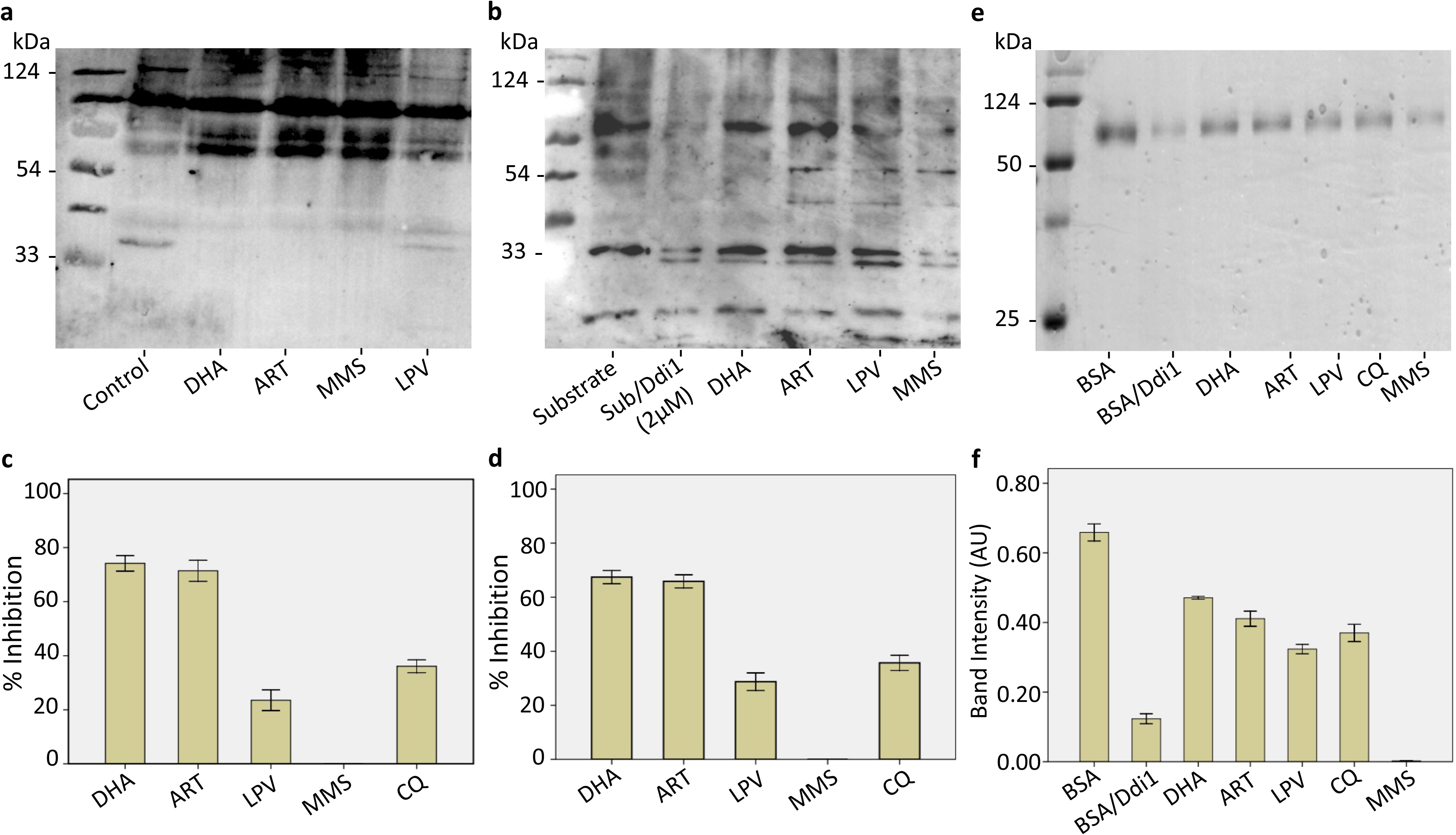
Artemisinin exposure enhances protein ubiquitination and blocks the *Pf*Ddi1 activity. (**a**) Western blot analysis showing increased ubiquitination in drug-treated *P. falciparum* parasites compared to the control. (**b**) Artemisinin (1 μM) blocked the cleavage of the polyubiquitin substrates. The parasite lysates were probed with rabbit anti-ubiquitin antibodies and the blots are a representative of three independent assays. (**c, d**) Percentage inhibition of the *Pf*Ddi1 enzyme activity against the retropepsin (c) and proteasome (d) substrate, determined at 3 hours (**e**) Coomassie stained SDS-PAGE showing inhibition of BSA degradation by *Pf*Ddi1. (**f**) Band intensity values of the inhibition of PfDdi1-catalyzed BSA degradation by the compounds. The intensity values are represented as arbitrary units (AU). The bars show means ± standard error for three independent reactions.

### Artemisinin treatment leads to increased recruitment of *Pf*Ddi1 into the nucleus following DNA damage

Artemisinin has been shown to induce DNA damage in malaria parasites as demonstrated by comet assays^9^. However, data on the nature of the DNA damage remains elusive. Using the *in situ* DNA fragmentation (TUNEL) assay, we observed DNA fragmentation (direct TdT-mediated dUTP nick end labeling) in more than 90% of the *P. falciparum* parasites following a two-hour exposure to artemisinin (Fig 3a). To gain insights into the possible molecular events accompanying the artemisinin-specific DNA fragmentation, we employed immunofluorescence assays (IFA), using anti-*Pf*Ddi1 antibodies, to evaluate the expression profile of the *Pf*Ddi1 protein, under drug pressure. Previously, Ddi1 was shown to repair DNA-protein crosslinks (DPCs) in yeast cells^28^. Our data showed that, whereas *Pf*Ddi1 is predominantly expressed in the cytoplasm, artemisinin and DHA treatment led to increased recruitment of the protein into the *P. falciparum* nucleus (Fig. 3b and c). Thus, whereas we have not shown the exact DNA repair mechanism, the shift in expression is likely to be a causal relation between the DNA fragmentation and *Pf*Ddi1 repair strategies.

**Fig. 3:**
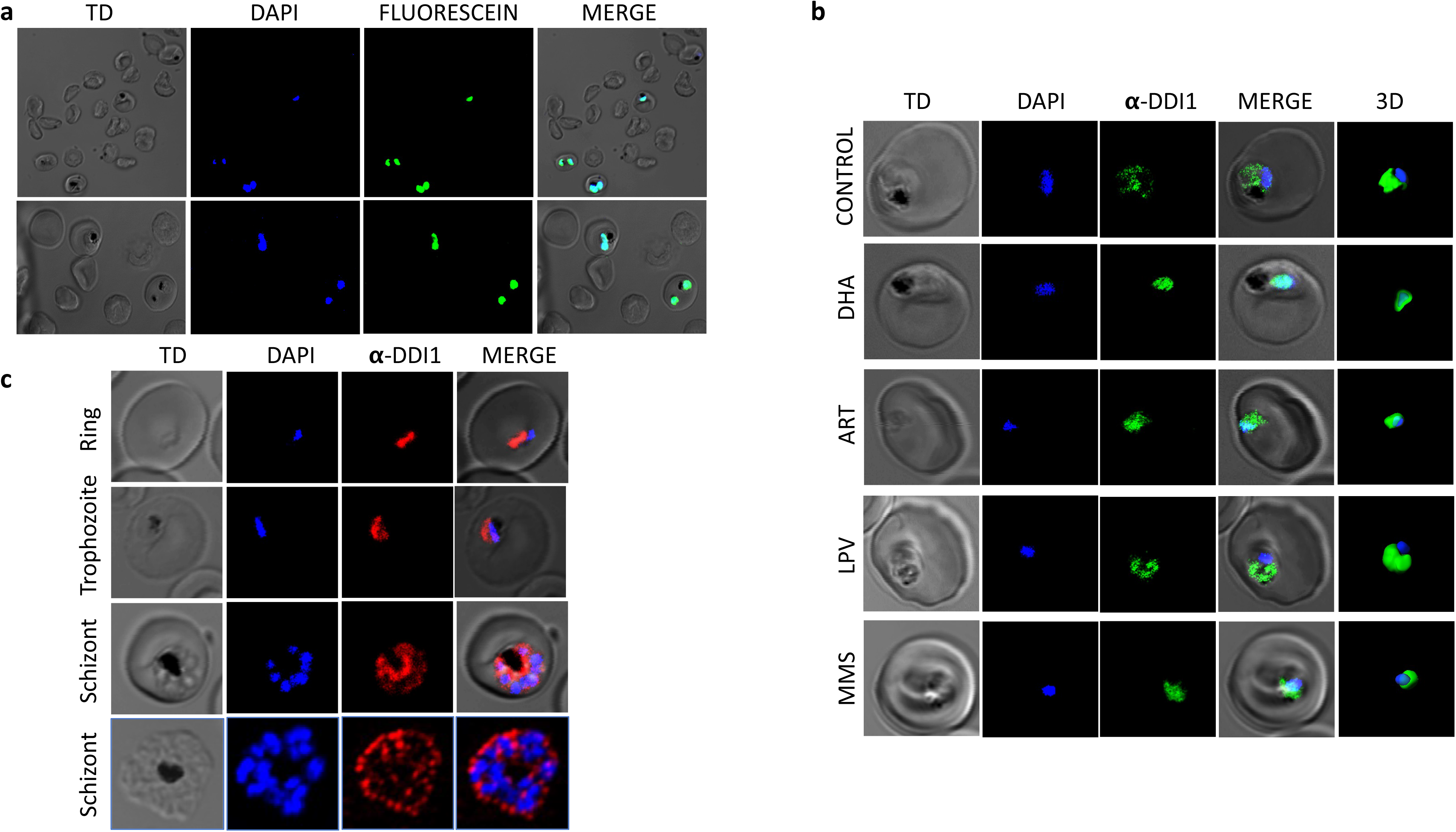
Causal relation between artemisinin-specific DNA fragmentation and *Pf*Ddi1 subcellular localization. (**a**) Representative IFA images showing artemisinin induces DNA fragmentation in P*. falciparum* parasites. The parasites were subjected to drug pressure for 1hr and then assayed using TdT-mediated dUTP nick end labelling. Fragmentation was observed in more than 95% of the infected RBCs. (**b**) Increased recruitment of *Pf*Ddi1 into the nucleus following artemisinin pressure compared with control (DMSO). The IFA staining was performed using mice anti-*PfDdi*1 antibodies and then underwent 3D reconstruction in IMARIS software. (**c**) IFA staining of *P. falciparum* blood stage parasites with rabbit anti-*Pf*Ddi1 antibody showing constant expression and localization of Ddi1 in the cytoplasm. The individual stains, merged images, and bright field are shown. Scale bars: 2 μm. The experiments were performed on 2–4 independent occasions with technical duplicates.

### Artemisinin binds and stably interacts with the highly conserved aspartic protease motif “DSG” in *Pf*Ddi1 protein

Having established the effect of artemisinin on the *Pf*Ddi1 protease activity, we carried out surface plasmon resonance (SPR) and Bio-Layer Interferometry (BLI) based binding assays, as well as computational analysis to delineate the exact interaction between *Pf*Ddi1 and artemisinin. Briefly, over 8500 response units (RU) or up to a maximum of 0.8nm shift of the recombinant *Pf*DdI1was immobilized via the amine coupling chemistry (CM5 chip) or streptavidin-biotin capture (Octet biosensors), respectively. Depending on the buffer in which the compounds were dissolved, we used either HBS-EP or DMSO as running buffer. Both artemisinin and MMS showed high affinity interactions with *Pf*Ddi1 in both bindig assays with k_D_ values of 1.06×10^−6^/1.56×10^−6^/, and1.70×10^−6^/2.51×10^−5^ respectively, while lopinavir showed lower binding affinities with a k_D_ value 2.22×10^−4^/5.62×10^−4^, in SPRand BLI assays (Fig. 4a-c). This binding was specific as none of the compounds showed interaction with the heme detoxification protein (HDP) (Supplementary fig. 5). On the other hand, PFAM and INTERPROSCAN search revealed the presence of two conserved domains in the *Pf*Ddi1 sequence: N-terminal Ubiquitin-like domain (4-74aa) and a retroviral-like protease domain (RVP; 222-345aa) (Supplementary Fig. 1a and 5). Further, to know conservation among different species, we performed multiple sequence alignment of Ddi1 sequences from *P. falciparum, L. major,* yeast and human. The alignment analysis showed that *Pf*Ddi1 had 95% query coverage and ~29% identity with human Ddi1 (hDdi1). In addition, all the aligned Ddi1 protein sequences showed higher conservation in the central RVP domain region as compared to the N- or C- terminal regions, with the presence of superimposed highly conserved aspartyl protease signature motif “D(S/T)G” (Supplementary Fig. 6).

**Fig. 4:**
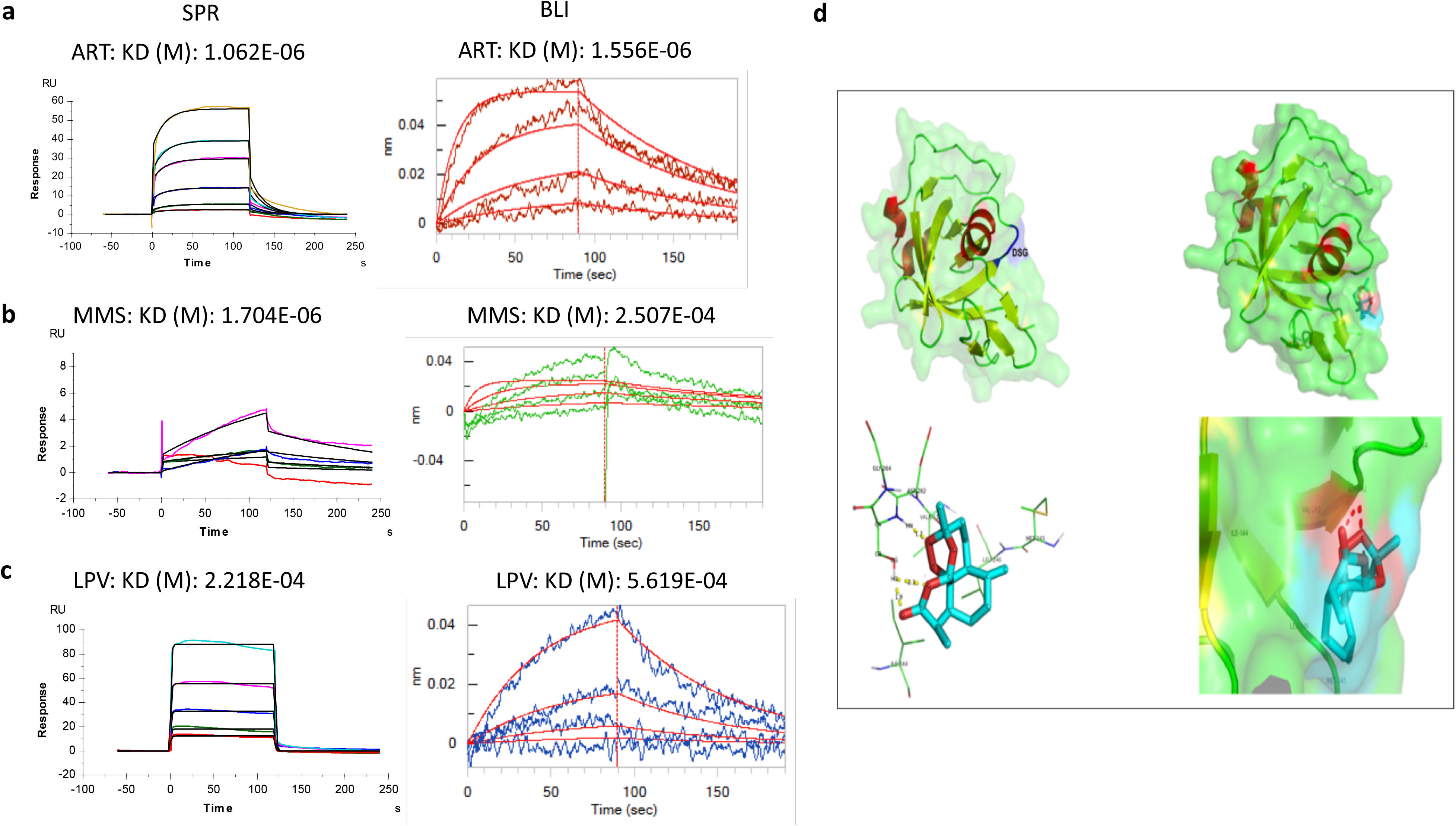
Artemisinin binds to the recombinant *Pf*Ddi1 protein: SPR, BLI and *In silico* assays. Binding of artemisinin (ART; **a**) and Methylmethanesulfonate (MMS; **b**) showed high affinity interactions with *Pf*Ddi1, compared to Lopinavir (LPV; **c**), as shown by the K_D_ values. Two independent SPR or BLI experiments were performed and representative binding sensorgrams are presented. Both the real time binding curves and the global 1:1 fits are shown. The binding kinetics data was analyzed using the Biacore T200 evaluation software v3.1 or the Octet Software v10.0. (**c**): Artemisinin shows stable interaction with the highly conserved aspartic protease motif “DSG” in *Pf*Ddi1 protein. PfDdi1 residues present within 4Å of artemisinin and is involved in direct interaction. Homology based 3D model of the PfDdi1 RVP domain as generated by SWISSMODEL. The conserved Aspartic protease motif DSG is present in the coil region. Red – helix, green – coil, yellow – sheet and blue – conserved active motif.

As no crystal structure for PfDdi1 protein is available so far, we generated a homology-based 3D model for *Pf*Ddi1. All attempts to generate complete stable 3D structure for *Pf*Ddi1 (382aa) were futile. We then generated a partial 3D model for the *Pf*Ddi1 RVP domain (243-366aa) using 4RGH, a human Ddi1 Homolog 2 protein, having 37% query coverage and 48.61% identity with the *Pf*Ddi1, as a template (Fig. 4b). The Ramachandran plot for the predicted model showed no residues in the disallowed region, confirming the good quality of the model. To assess whether an artemisinin molecule binds to the protease domain region, we performed *in silico* docking using AutoDocktools. Here, site specific docking was performed using the predicted *Pf*Ddi1 as a receptor and an artemisinin molecule as a ligand. Grid box was generated using nitrogen of Asp262 as the center (grid points xyz coordinates as 40, 40, 40 and spacing of 0.4Å), and the other default parameters were used for the screening. The docking analysis revealed *Pf*Ddi1 protein binding with artemisinin in the active catalytic protease signature motif (DSG) (Fig. 4d). The free binding energy for the reaction was −5.81 kcal/mol. Hydrogen bonds were formed between artemisinin and Ser263 (in the catalytic DSG motif), with all the three active-site residues of the aspartyl proteases present within 4Å of artemisinin. Taken together, the binding and docking results suggest stable interaction between the *Pf*Ddi1 protein and artemisinin, and possible inhibition of the *Pf*Ddi1 protease activity by artemisinin.

### *Pf*Ddi1 restores the protein secretion phenotype in yeast cells

To know whether the *Pf*Ddi1 is a true orthologue of yeast Ddi1, we performed complementation studies in *S. cerevasie* yeast cells. We singly disrupted the *Sc*Ddi1 gene, by homologous recombination, and assayed whether the *Pf*Ddi1 ortholog could complement the phenotypes in the knockout yeast cells. Cells bearing a *Ddi1* gene disruption were seen to grow normally. However, as has been shown previously^29^, our data showed that *Ddi1*Δ yeast cells secreted significantly higher protein levels into the media (Fig. 5a). On average, *Ddi1*Δ yeast cells secreted more than ~30% of proteins into the media, compared with the wild type strain. To test whether *Pf*DdI1 restores the wild-type proteins secretion phenotype, we cloned a gene encoding the full-length *Pf*Ddi1 into a yeast expression vector pGPD2. The ligated construct was transformed into the *DdI1*Δ yeast cells. The *Pf*DdI1 construct was able to revert the protein secretion phenotype to WT level, i.e the level of protein secretion decreased in comparison to the knock-out strain (Fig. 5a).

**Fig 5:**
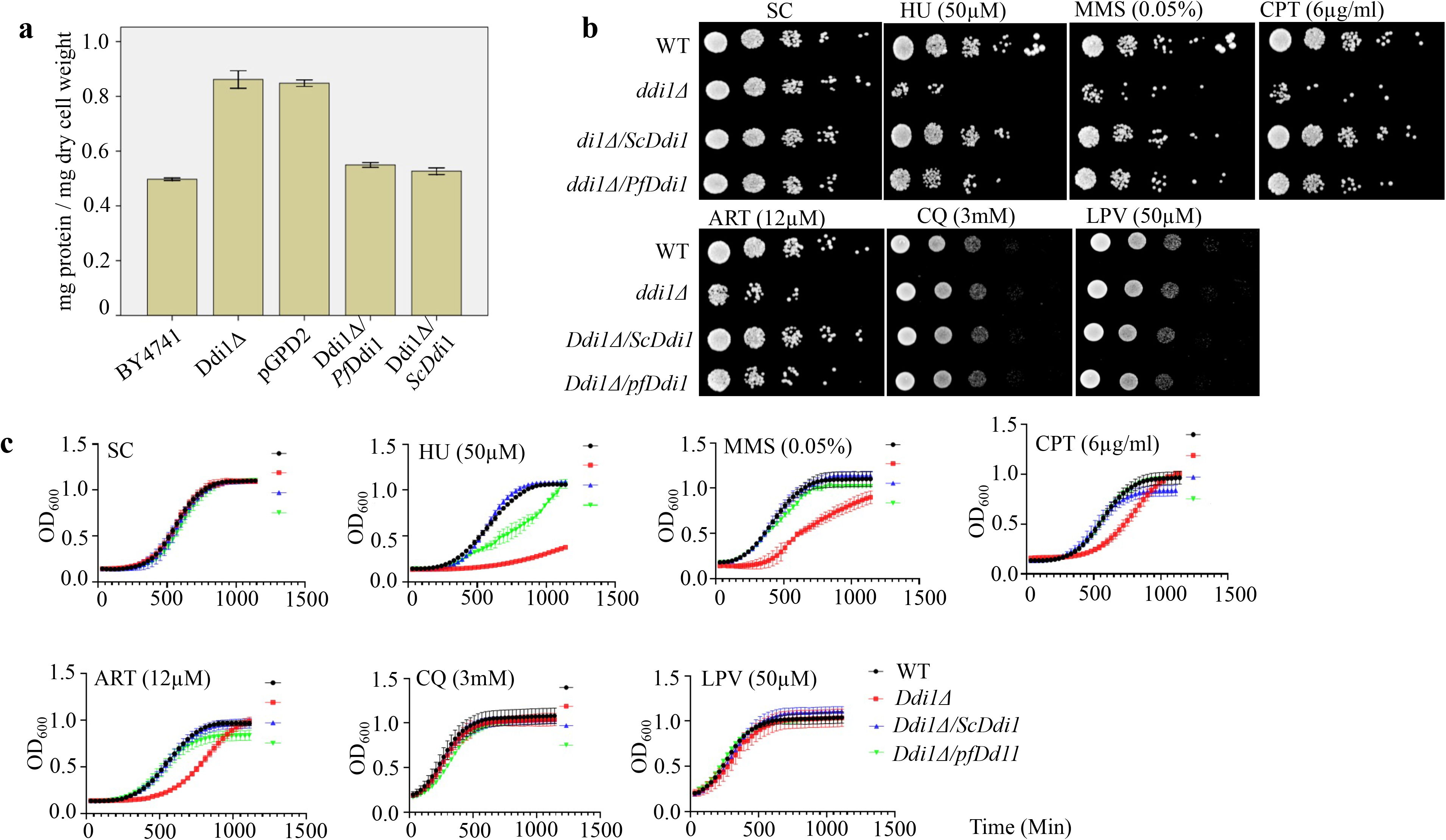
Ddi1 deficient yeast cells are more susceptible to artemisinin pressure. **(a)** Ddi1 deficient yeast cells secret high levels of proteins into the media and *Pf*Ddi1 reverts the protein secretion phenotype to wild type, as did the *Sc*Ddi1 construct. The protein content in the supernatant was estimated using the Pierce BCA Protein Assay Kit and the protein concentration expressed as milligrams of protein secreted per milligram of dry cell weight. Triplicate assays were conducted, and the bars represent means ± standard error. Spot test images (**b**) and growth curve assays (**c**) showing the effect of the compounds on the different yeast lines. Whereas deletion of Ddi1 did not affect the growth fitness of the yeast cells, Ddi1 deficient cells were more susceptible to artemisinin.

### Ddi1 deficient yeast cells are more susceptible to Artemisinin pressure

It had been previously shown that mutations in some DNA repair genes confer resistance to Artemisinin^30^. Moreover, DNA damaging agents have been shown to perturb and induce transcriptional changes in 21% of the *P. falciparum* genome^10^. These changes involve up-regulation of the genes of the DNA repair machinery. Similarly, yeast studies demonstrated that DNA damaging agents trigger differential expression in one third of the entire *S. cerevisiae’s* gene pool^31^. We reasoned that since the proteasome is central to the repair or disposal of damaged cellular components, the yeast cells lacking the Ddi1 might be more susceptible to DNA damaging agents. We, therefore, incubated equal amounts of yeast cells with different drugs and DNA Damaging agents such as artemisinin, chloroquine, lopinavir, hydroxyurea, MMS, and camptothecin and measured sensitivity using both OD (growth curves) and spotting tests. Our results demonstrated that Ddi1Δ yeast cells were more susceptible to artemisinin (12μM). These Ddi1Δ cells were also hypersensitive to DNA damage drugs; hydroxyurea, MMS, and camptothecin (Fig.5b and c). Together, these results augments our observations in *P. falciparum* and demonstrate that *Pf*Ddi1 reduces the sensitivity of the cells to artemisinin insults, and artemisinin works similar to the known DNA dmaging agents.

## Discussion

The spectrum of drugs to which the human malaria parasite, *Plasmodium falciparum* has not evolved tolerance is rapidly diminishing. Reports on decreased sensitivity of the parasites towards the recommended first-line treatment for *P. falciparum* malaria, artemisinin, threatens the global efforts to combat the disease. To circumvent the resistance, improve the efficacy or generate new drugs, it’s critical to understand the mechanisms of action/resistance of the artemisinin. Since it has been shown that artemisinin causes indiscriminate damage to parasite cellular proteins^7–11^, recruitment of the parasite protein repair machinery might be pivotal in assuring parasite growth fitness. Besides, artemisinin compromises the functions of the parasite proteasome and synergizes with proteasome inhibitors in the killing of artemisinin resistant parasites^12,18^. Here, we investigate the mode of action of artemisinin on the parasite proteasome machinery. Expression and activity analysis of *Pf*Ddi1, a proteasome shuttle protein with an unusual RVP domain, show that *Pf*Ddi1 is an active A_2_ aspartyl protease that hydrolyzes proteasome substrates, including polyubiquitin proteins. However, the enzyme could not catalyze the hydrolysis of Bz-RGFFP-MNA, a cathepsin D substrate. This activity is in line with earlier reports that showed *L. major* Ddi1 as an active aspartyl proteinase^26^, as well as *Sc*Ddi1 as a ubiquitin-dependent protease that acts on polyubiquitinated substrates^23^. The ability of *Pf*Ddi1 to cleave polyubiquitinated substrates suggests that the *Pf*Ddi1 enzyme might not only be a shuttle protein but could inherently degrade damaged proteins. Thus, the *Pf*Ddi1 protein might be acting synergistically with the proteasome machinery to degrade the ubiquitinated proteins. Indeed, previous reports have demonstrated that Ddi1 compensates for proteasome dysfunction in *Caenorhabditis elegans*^22^. In addition, deletion of Ddi1 (ΔDdi1) in *T. gondii* results into accumulation of ubiquitinated proteins, a phenomenon ehnanced by double deletion (ΔDdi1 and ΔRad23)^24^. Therefore, the essentiality of the *Pf*Ddi1 advocates its multiple roles in parasite life cycle and negates any redundancy in its functions. On the other hand, *in silico* data reveals that unlike most of the Ddi1 analogs, *Pf*Ddi1 lacks the UBA domain, thus suggesting that the UBA domain does not contribute to the protease activity of the Ddi1 protein. These results are consistent with the observation made earlier for *Sc*Ddi1, which shows that the deletion of UBA domain had no effect on the activity of *Sc*Ddi1^23^.

To further show that *Pf*Ddi1 is a functional homolog of the *Sc*Ddi1, which is one of the best characterized Ddi1 protein, complementation studies were carried out in *S. cerevisiae* cells. Functional expression of the *Pf*Ddi1 in *S. cerevisiae* cells showed its ability to restore disrupted phenotypes. We show that, unlike in *P.falciparum* where functional disruption of Ddi1 gene is deleterious, *Sc*Ddi1 gene is not refractory to deletion in yeast cells. However, as previously reported^29^, deletion of *Sc*Ddi1 increased secretion of proteins to the growth media. Interestingly, despite the differences in the domain structure, the *Pf*Ddi1 gene robustly complemented the yeast secretion phenotype. This observation might infer that the C-terminal UBA domain lacks crucial sequences associated with the suppression of protein secretion.

Since artemisinin has been shown to promiscuously target parasite proteins and induce DNA damage (Comet assay), we sought to define whether it also inhibits the activity of *Pf*Ddi1. We first demonstrate that, indeed, artemisinin causes protein damage which leads to piling up of polyubiquitinated proteins. This is in agreement with previous data which showed that the artemisinins induce polyubiquitination in the malaria parasite^12,16^. On the other hand, we demonstrate that artemisinin causes DNA damage by directly inducing DNA fragmentation in the *P. falciparum* parasites. Interestingly, the exposure of the parasites to genotoxic artemisinin insults causes increased recruitment of *Pf*Ddi1 into the nucleus. This suggests that *Pf*Ddi11 might be involved in the regulation of DNA damage response to artemisinin. Indeed, previous reports have implicated Ddi1 in the repair of DNA-protein crosslinks ^28^. The exact mechanism adopted by the *Pf*Ddi1 in the DNA damage repair in the parasites remains of utmost interest. Enzyme inhibition assays showed that artemisinin blocked 71.4% or 65.9% of the activity of *Pf*Ddi1 against the retropepsin or proteasome substrates, respectively. Besides, artemisinin significantly inhibited the degradation of polyubiquitinated substrates, a finding that unequivocally fortifies the inhibitory effect of artemisinin on the *Pf*Ddi1activities. However, lopinavir (50 μM), a known HIV protease inhibitor, could only yield marginal inhibition (~23.5%) of the activity of *Pf*Ddi1, a retropepsin protease. In addition, interaction sensograms from binding assays and *in silico* modeling and docking studies showed high affinity binding between artemisinin and *Pf*Ddi1. To provide additional evidence on the role of Ddi1 in the mediation of artemisinin activities, we studied the growth fitness of Ddi1 deficient *S. cerevisiae* (*Sc*DdI1Δ) cells. This transgenic line showed differential susceptibilities to an array of DNA damage compounds as well as to artemisisnin, the mainstay anti-malarial drug. *Sc*DdI1Δ cells were more suceptible to artemisinin pressure, compared to the wild type cells. Therefore, the observed increased susceptibility might be as a result of the *Sc*DdI1Δ cells’ inefficiency to invoke DNA and protein repair upon artemisinin-induced damage. In fact, artemisinin has been shown to increase the generation of free radicals that chokes the yeast cell homeostasis^32^. Besides, artemisinin has been shown to elicit DNA damaging effect comparable to MMS, an alkylating agent^10^. Restoration of the WT growth fitness by the *Pf*Ddi1 infers that the UBA domain plays an insignificant role in responding to artemisinin genotoxic insults.

These results thus enhance the evidence on the mode of action of artemisinin that has been earlier shown to kill the parasites via a two-step mechanism; causing ubiquitous protein damage and compromising parasite proteasome functions^12^. Therefore, based on our data coupled with the previous observations, we propose that artemisinin exerts its pressure on the parasite by compromising the *Pf*Ddi1 protein, an important player in the parasite protein homeostasis. The compromised *Pf*Ddi1 does not only lose its ability to degrade the damaged proteins but also curtail its shuttling capacity, thus leading to accumulation of the damaged proteins and eventual death of the malaria parasite (Fig. 6).

**Fig. 6:**
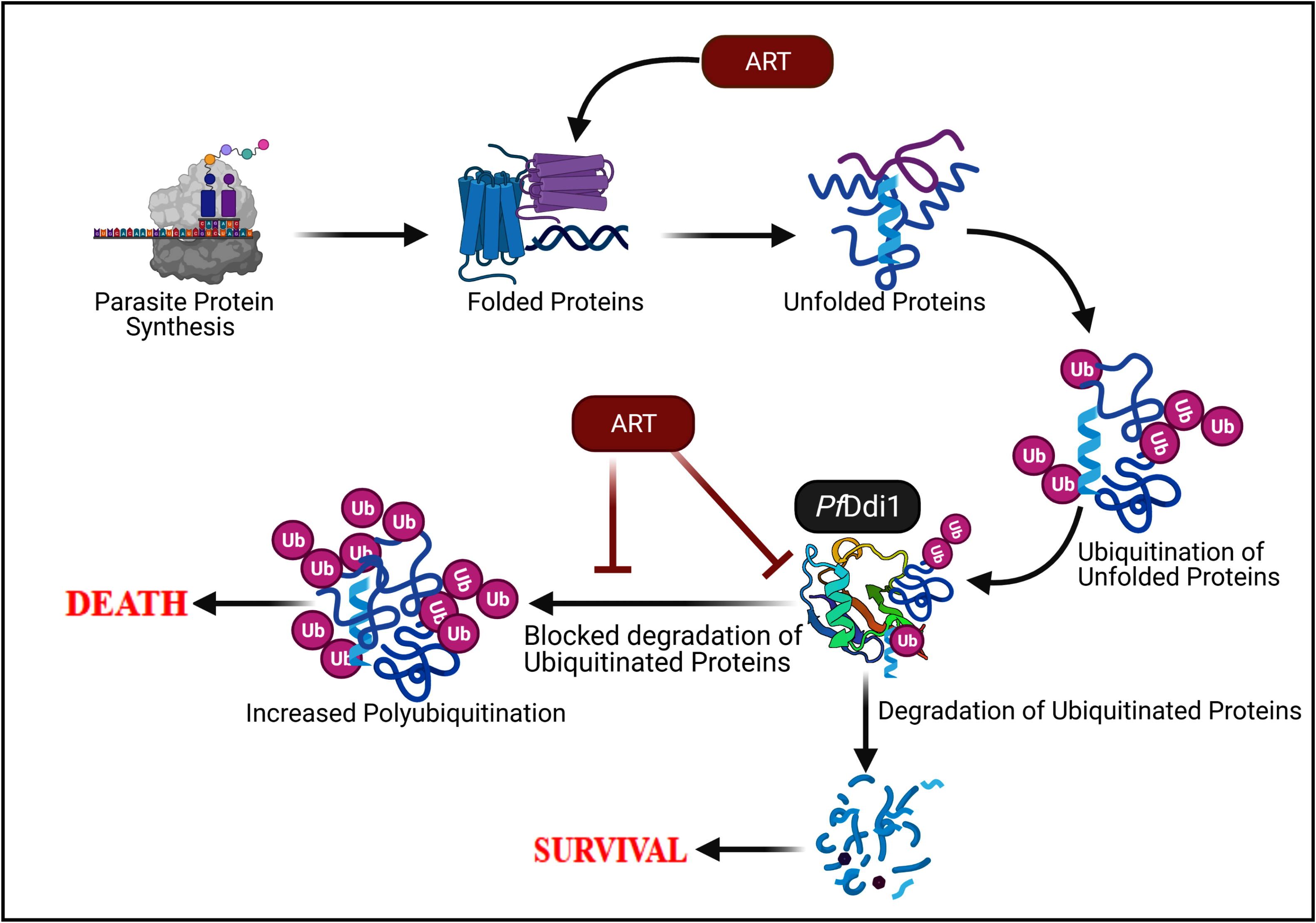
Model demonstrating ART-induced killing of *Plasmodium* parasites. ART has been shown to target several parasite proteins and processes. Here, we focus on the role of the *Pf*Ddi1 in mediating the actions of ART. The ART’s ubiquitous damage of parasite proteins leads to the need for tidying up via the *Pf*Ddi1 or proteasome machinery. Besides causing the protein damage, artemisinin binds to *Pf*Ddi1 and blocks the degradation of the damaged proteins. Besides, ART might be preventing the trafficking of the damaged proteins to the proteasome for degradation. The blockage leads to accumulation of the damaged proteins, choking the parasites thus leading to the ultimate death.

In conclusion, here we show that *Pf*Ddi1, an essential *P. falciparum* protein, is an active A_2_ family aspartic protease, with inherent abilities to degrade polyubiquitinated proteins. *Pf*Ddi1 is a true orthologue of *Sc*Ddi1 and could be involved in DNA damage repair strategies by the parasites. We further show that artemisinin, a first line drug against *P. falciparum* malaria, kills the parasites by inducing protein damage and inhibiting tidying up by blocking the activity of *Pf*Ddi1, a unique ubiquitin-proteasome retropepsin, that results in piling up of the damaged proteins. These results thus provide insight into the mode of action of artemisinin and pave the way for development of new antimalarial drugs targeting *Pf*Ddi1.

## Methods

### Ethics statement

All protocols were conducted in accordance with prior approvals obtained from the International Centre for Genetic Engineering and Biotechnology (ICGEB)’s Scientific Ethical Review Unit and the Institutional Animal Ethics Committee (IAEC; ICGEB/IAEC/02042019/MB-7).

### Cloning, expression, and purification of recombinantPfDdi1

The *Pf*DdI1 gene (PF3D7_1409300) was amplifiedfrom genomic DNA using specific primers; DdI1 forward 5’ -GCGGATCCATGGATATGGTTTTTATTACAATATCAGACG-3’, reverse 5’ GCGTCGACCTCGAGTAAATCATTGTTTGCATCAATG–3’. The PCRproduct was first cloned intopJET vector (Thermo Scientific) and then sub-cloned into pET-28b expression vector using NcoI and XhoI restriction sites (Thermo Scientific). The pET-28b clone was expressed in Rosetta (DE3) *Escherichia coli* cells (Invitrogen). The cells were grown to mid log phase, and then induced with isopropyl-1-thio-β-D galactopyranoside (IPTG, 1 mM) for 14 h at 16°C. The bacterial culture was harvested by centrifugation at 4000xg for 20 min. The cell pellet was re-suspended in lysisbuffer (50 mMTris·HCl at pH 8.0, 200 mMNaCl, 1.0% Triton X-100 and 1.0% PMSF) and then sonicated.The supernatant was collected by centrifugationat 9,000 rpm for 50 min, at 4°C. The soluble recombinant protein was purified using theNickel-Nitrilotriacetic acid (Ni-NTA^+^; Qiagen) resin. Briefly, the protein was allowed to bind in 20 mM imidazole-containing binding buffer (50 mMTris:HCl at pH 8.0 and 200 mMNaCl) for 3 h at 4 °C. The resin with bound protein was washed in 30 mMimidazole-containing binding buffer and then the bound protein was eluted in varying concentrations of the imidazole (50, 75, 100, 150, 200, 300 and 500 mM) in 50 mMTris:HCl at pH 8.0 and 200 mMNaCl. The purified fractions were checked on SDS-PAGE and Western blot analysis using α-His antibodies. All the pure fractions were pooled and dialyzed in the Tris-Nacl buffer (50 mMTris-Hcl pH 8, 200 mMNacl), and then concentrated.

### Generation of antibodies against *Pf*Ddi1

All animal protocols were conducted in accordance with prior approvals obtained from the International Centre for Genetic Engineering and Biotechnology (ICGEB)’s scientific review committee and the institutional animal ethics committee (ICGEB/IAEC/02042019/MB-7). We used BALB/c inbred mice and female NZW rabbits to raise anti-bodies against the recombinant *Pf*Ddi1. The mice were immunized with 20μg of the protein while the rabbits were immunized with 200μg protein in the presence of complete/incomplete Freund’s adjuvant, using the i.p. and s.c. modes of injection, respectively. After the third bleed, the antibody titers were quantified by ELISA. The specificity of the raised antibodies was analyzed on the recombinant *Pf*Ddi1 protein and the *P. falciparum* parasite lysate.

### *Pf*Ddi1 enzymatic assays

The aspartyl proteinase activity of the purified recombinant *Pf*DdI1 was probed against three substrates; Bz-RGFFP-4MβNA, DABCYL-Gaba-SQNYPIVQ-EDANS orSuc-LLVY-AMC (Bachem, Bubendorf, Switzerland), following the protocol described earlier ^26^. Briefly, 2.0 μMof the recombinant *Pf*DdI1was incubated with decreasing concentration (80 μM-1.25 μM) of each of the substrates in 100 mM sodium acetate buffer, at pH 5.0. Triplicate assays were carried out in a total volume of 200 μl in 96 well opaque plates, for 4 h at 37 °C. Assays with Heme Detoxification Protein (*Pf*HDP) were used as the control. Both the DABCYL-Gaba-SQNYPIVQ-EDANSand Suc-LLVY-AMC cleavage signals were measured at an excitation wavelength of 355nm and an emission wavelength of 460nm. On the other hand, an excitation and emission wavelength of 340 and 425 nm, respectively, was used to monitor the hydrolysis of Bz-RGFFP-4MβNA. The fluorescence signals were captured at 15-minute intervals with the VICTOR Multilabel plate reader (VICTOR X3). Due to the intrinsic reduction of fluorescence associated with fluorescence resonance energy transfer (FRET)-based cleavage assays, fluorescence from varied concentrations of free Edans (from 0.625 to 40 μM) in the assay buffer was used to generate a standard calibration curve and for correction of the inner filter effect^33,34^. The obtained relative fluorescence units were converted into velocity {μg (cleaved substrate)/s} and then used to derive the kinetic and catalytic constants in GraphPad Prism v6.0. The enzyme’s overall ability to cleave the substrate was represented ask_cat_/K_M_ (M^−1^ s^−1^).

### Proteolytic assays on polyubiquitin substrates and macromolecules

We incubated 20μg of polyubiquitin substrate (K^48^-linked) or 0.25mg/mL of bovine serum albumin with 2.0 μM of freshly purified recombinant *Pf*Ddi1in the 50 mM sodium acetate buffer (as described previously), pH 5.0, in a final volume of 100 μl. Triplicate assays and the control (substrate alone) mixture were incubated at 37°C for 2h. The mixtures were centrifuged and then resolved in a 12% SDS-PAGE. Cleavage of the polyubiquitin substrate was probed using rabbit anti-ubiquitin antibodies and then detected by enhanced chemiluminescence (ECL) using the Bio-Rad ChemiDocTM MP imaging. On the other hand, cleavage of the BSA substrate was stained by coomassie brilliant blue. The arbitrary band intensity values were presented as means ± standard error (SE).

### In vitro culture of *Plasmodium falciparum* and drug treatment

*P. falciparum* parasites (3D7 strain) were cultured and maintained in purified human red blood cells at 4% hematocrit, in RPMI 1640 medium (Gibco) supplemented with 0.25% Albumax I (Gibco), 2 g/L Sodium bicarbonate (Sigma), 0.1 mM hypoxanthine (Sigma), and gentamicin (Gibco). Parasite cultures were kept at 37°C with 5% CO_2_, 3% O_2_, and 92% N_2_. The parasites (ring stage; 2-4 hpi) were tightly synchronized with 5% (v/v) D-sorbitol (Sigma) and then monitored by Giemsa staining of methanol-fixed blood smears. Tightly syncronised mid-trophozoites were diluted to 5% parasitaemia and then subjected to the drug treatment (artemisinin; 1μM, DHA; 1μM, MMS; 0.05% or LPV; 50μM, for 4hr. DMSO was used as a vehicle treatment for all the assays. The parasite cell pellets were washed with ice-cold PBS and then lysed with 0.15% (w/v) saponin and radioimmunoprecipitation assay buffer (RIPA buffer) as described previously. The protein content was normalized with BCA assay and then resolved by a 10% SDS PAGE. The gel was transferred to a nitrocellulose membrane blocked with 5% (w/v) skim milk for 1 h at room temperature and probed with primary rabbit anti-ubiquitin antibody (1:100) overnight at 4 °C, followed by HRP-conjugated secondary antibody for 1 h at room temperature. The blots were processed by ECL reagents and then detected using the Bio-Rad ChemiDocTM MP imaging.

### Enzyme inhibition assays

For the enzyme inhibition assays, we preincubated 2.0μM of the enzyme with drug compounds { artemisinin (1μM), DHA (1μM) MMS (0.05%) or LPV (50μM), in sodium acetate buffer, pH 5.0 for 10 minutes, at ~24°C. We then added 10 μM of the fluorescence substrates or 20μg of polyubiquitin protein and the inhibition experiments were carried out at 37°C. The fluorescence signals and the protein degradation were processed as early described. The experiments were carried out in triplicates and fluorescence inhibition was expressed as a percentage of the control.

### Protein-drug interaction assays (optical methods; SPR and BLI)

All the SPR or BLI experiments were performed using a T200 instrument (Biacore) or the Bio-Layer Interferometry (BLI) Octet RED96e platform (FortéBio). Freshly prepared HEPES buffered saline (HBS)-EP (0.01 M HEPES; pH 7.4, 0.15 M NaCl, 0.003 M EDTA, 0.05% vol/vol P20 surfactant) or DMSO was used as running buffer for the experiments. For the SPR interaction assay, over 8500 response units (RU) of the recombinant *Pf*DdI1in sodium acetate buffer (pH 4.5) was immobilized on a SPR CM5 sensor via amine coupling^35^. A blank flow cell was used for reference corrections. Heme detoxification protein (HDP) was also immobilized on the CM5 sensor and used as the control protein.For the BLI interaction assay, biotinylated*Pf*Ddi1, diluted to a concentration of 25 μg/mLin kinetics buffer (HBS-EP with 0.1 mg/ml BSA) was immobilized on streptavidin-coated (SA) biosensors (FortéBio).The ligand was immobilized up to a maximum of 0.8 nm shift. Reference biosensors loaded with the ligand but dipped into wells containing only the buffer were run in parallel to control for possible drifts and establishment of baseline. Serial two-fold dilutions of the compounds; artemisinin (1μM), lopinavir (50μM) or MMS (0.05%), diluted in the running buffer, were used and the kinetics performed at 25°C. In the SPR experiments, a total of 0.2mL of the sample was injected while in BLI, each biosensor was stirred in 0.2 mL of the sample at 1000 rpm. The kinetics data was analyzed using the Biacore T200 evaluation software v3.1 or the Octet Software v10.0. The affinity between the immobilized protein and the compounds was expressed as dissociation constant (K_D_).

### *Pf*Ddi 3D model generation and *in silico* docking

PlasmoDB(https://plasmodb.org/plasmo/) andUniProt(https://www.uniprot.org/) were used to retrieve sequences for Ddi1 proteins^36^.Artemisinin chemical structure was retrieved from PubChem database (https://pubchem.ncbi.nlm.nih.gov/). To identify conserved domains in the *Pf*Ddi1 protein, we performed PFAM and INTERPROSCAN search^37,38^. We then used CLUSTAL-Omega version 1.2.4 to perform multiple sequence alignment^39^. SWISS-model was used to generate three dimensional (3D) model for *Pf*Ddi1^40^, and then Rampage tool was used for quality check analysis for the predicted model^41^. AutoDock tools were used to perform docking analysis^42^ and the generated images of the molecular models were visualized using PyMol (https://pymol.org/2/).

### Immunofluorescence assay

Immunofluorescence assay (IFA) was performed with *Plasmodium* parasite cells in suspension. Briefly, parasite pellet was washed in 1×PBS and fixed in 4% v/v formaldehyde supplemented with 0.0075% v/v glutaraldehyde in PBS for 30 min at RT. The cells were then permeabilized in 0.1% Triton X-100 in PBS for 20 min and then washed in 1×PBS. We then blocked with 5% BSA for 1h at room temperature, and then incubated with primary antibodies (anti-Ddi1) overnight at 4°C, followed by incubation with fluorophore-conjugated secondary antibodies (1:100,000 in 3% BSA). DAPI was added and incubated for 20 min. Thin blood smears of the stained cells were made on microscope slides and mounted with cover slips. The slides were imaged using a Nikon Eclipse Ti-E microscope. Images were processed using the NIS-Elements AR (4.40 version) software. For 3D reconstruction, we used Imaris x64 version 6.7 (Bitplane).

### *In Situ* DNA Fragmentation (TUNEL) Assay

Treated or solvent alone cells (trophozoite-rich) were fixed and permeabilized as described above. DNA fragmentation was assessed by TUNEL using *In Situ* Cell Death Detection Kit, TMR Red (Roche Applied Science, Mannheim, Germany), following the manufacturer’s instructions. Briefly, the permeabilized cells were incubated with TdT enzyme and fluorescein-12-dUTP for 1 h at 37°C, followed by DAPI for 20 min and then washed in 1 × PBS. Thin blood smears of the labeled parasite cells were made on microscope slides and then imaged as described above. The percentage of TUNEL-positive cells was estimated.

### Generation of transgenic yeast cells

To know whether the *Pf*Ddi1 is a true yeast orthologue, we performed complementation studies in *S. cerevasie* yeast cells. We amplified deletion constructs using primers bearing the nourseothricin (NAT) selection marker (Supplementary Table 1). The genes were deleted by homologous recombination and the integration confirmed by PCR-based genotype analysis. On the other hand, the genes encoding full-length *Pf*Ddi1 were amplified from gDNA using primers as shown in supplementary Table 1. The PCR products were cloned into pGPD2 yeast expression vector at SpeI/XhoI site. The constructs were then transformed into the the *S. cerevisiae* strain BY4741 by the lithium acetate method^43^.Selection of transformants were performed by plating over synthetic complete (SC) medium lacking uracil. Deletions were confirmed by genomic DNA PCR with appropriate set of primers (Supplementary Table 1).

### Phenotypic characterization

#### Protein secretion assay and growth rate

The secretion assay was performed following the protocol described by^29^. Briefly, we inoculated a single colony from each strain into 5mL synthetic complete (SC) medium (0.67% YNB with all amino acids but not uracil) in conical centrifuge tubes. The culture was incubated for 48h at 30°C in an orbital shaking incubator.We then pipetted 1 ml ofculture into pre-weighed microcentrifuge tubes and separated the supernatantfrom pellet by centrifugation for 5 min at 13 000 rpm. The protein content in the supernatant was estimated using the Pierce BCA Protein Assay Kit (Thermo Scientific) with BSA as a standard. The cell pellet was dried at 100°C and weighed.The assays were done in triplicates and the concentration was expressed as milligrams of protein secreted per milligram of dry cell weight.

#### Treatment with (genotoxic) compounds

Yeast cells were grown to mid log phase and adjusted to 0.1 OD_600_.Serial dilutions were prepared and spotted on SC-based agar plates supplemented withhydroxyurea (50μM), methylmethanesulfonate (0.05%), camptothecin (6μg/ml), artemisinin (12μM), chloroquine (3mM), or lopinavir (50μM). The plates were incubated at 30°C for 48 h. For yeast growth curve assays, we followed the protocol as described by^44^, with few modifications, in a liquid handling system (Tecan, Austria). Briefly,we inoculted a 96-well microplate with 5μL of fresh midlog-phase cell cultures. Each well contained SC media supplemented with either of the compounds in a total volume of 0.2 mL.The cells were incubated for 24 hours at 30°C and the cell population was recorded at an interval of 30 min. All the samples were prepared in triplicates.

### Statistical analyses

We exported the data to Excel (Microsoft) and carried out statistical analyses and data representation usingSPSS Statistics v16or GraphPad Prism v6.0. Nonlinear regression analysis was used to determine the enzyme kinetic constants (K_*m*_ and V_max_). The drug binding affinities (K_D_ values) were calculated after analysis of the association and dissociation from a 1:1 binding model. The results (bars) represent means ± standard error. The authors declare that they have no conflicts of interest with the contents of this article.

## Acknowledgement

I thank the International Centre for Genetic Engineering and Biotechnology (ICGEB) for awarding me the Arturo Falaschi ICGEB Predoctoral fellowship (F/KEN18-10) and providing the state-of-the-art facility at the ICGEB, New Delhi component, for the execution of my experiments. This work has been funded by JC Bose fellowship (DST/21/015) conferred to Dr Pawan Malhotra by SIBRI (Department of Science and Technology, India) and Flagship Grant given to ICGEB (DBT/10/026). I am grateful to Dr. Dinakar Salunke for critically reviewing the manuscript. I also thank Prof. Goldberg, Daniel (Washington University) for his enormous contribution and criticism in the development of this manuscript.

## Conflict of interest

The authors declare that they have no conflict of interest.

